# Translation control by maternal Nanog promotes oocyte maturation and early embryonic development

**DOI:** 10.1101/2022.08.16.504217

**Authors:** Mudan He, Shengbo Jiao, Ding Ye, Houpeng Wang, Yonghua Sun

## Abstract

Many maternal mRNAs are translationally repressed during oocyte maturation and spatio-temporally activated during early embryogenesis, which is critical for oocyte and early embryo development. By analyzing maternal mutants of *nanog* (M*nanog*) in zebrafish, we demonstrated that Nanog tightly controls translation of maternal mRNA during oocyte maturation via transcriptional repression of *eukaryotic translation elongation factor 1 alpha 1, like 2* (*eef1a1l2*). Loss of maternal Nanog led to defects of egg maturation, increased endoplasmic reticulum (ER) stress, and an activated unfold protein response (UPR), which was caused by elevated translational activity. We further demonstrated that Nanog, as a transcriptional repressor, represses the transcription of *eefl1a1l2* by directly binding to the *eef1a1l2* promoter during oocyte maturation. More importantly, depletion of *eef1a1l2* in *nanog* mutant females effectively rescued the elevated translational activity in oocytes, egg quality defects, and embryonic defects of M*nanog* embryos. Thus, our study demonstrates that maternal Nanog regulates oocyte maturation and early embryogenesis though translational control of maternal mRNA via a novel mechanism, in which Nanog acts as a transcriptional repressor to suppress transcription of *eef1a1l2*.

## Introduction

During oocyte development, lots of maternal mRNAs are transcribed and accumulated, and translational activation or repression of maternal mRNA in temporal and spatial determines oocyte development, maturation, and early embryogenesis. Translation of many maternal mRNAs is repressed during oocyte development (Evans and Hunter, 2005; Pique et al., 2008; Gosden and Lee, 2010), and maintenance of translational arrest of maternal mRNA is essential for normal oocyte development and maturation (Richter and Lasko, 2011; Yarunin et al., 2011). Failure of translational repression of maternal mRNAs resulted in various developmental defects, including apoptosis of oogenesis, impaired oocyte maturation and unsuccessful early embryonic development.

Various mechanisms of translational control have been described using different animal models. In zebrafish, the RNA-binding protein Zar1 binds to zona pellucida (ZP) mRNAs and represses translation of the ZP gene during oogenesis, loss of Zar1 induces oocyte apoptosis and ovary degeneration (Miao et al., 2017). An RNA-binding protein Ybx1 associates with processing body components and repress global translation activity during early embryogenesis (Sun et al., 2018). In *Drosophila*, translation of many germ cell-specific mRNAs are repressed by RNA-binding proteins during oocyte maturation. Germline RNA, *oskar* and *pgc*, are targeted by Bruno or Pumilio at the 3’-untranslated region (UTR) and translationally repressed during oogenesis, and mutation of RNA-binding sites results in precocious translation and mislocalization of mRNA (Kim-Ha et al., 1995; Snee et al., 2008; Flora et al., 2018). Me31B mediates the translational silencing of both maternal mRNAs during the maternal-to-zygotic transition (MZT) and oocyte-localizing RNAs in the transport process to oocyte (Nakamura et al., 2001; Wang et al., 2017). During the transport of *nanos* mRNA to oocyte, the failure of translational repression of *nanos* mRNA by Smaug would lead to ectopic translation of *nanos* and defects of anteroposterior axis formation (Dahanukar and Wharton, 1996; Smibert et al., 1996; Smibert et al., 1999). To date, the studies of translational repression in oogenesis have focused mainly on the posttranscriptional regulation, in which the translational repressors are mainly RNA-binding proteins. It remains elusive whether there is a general translational repressor that regulates the translation of maternal mRNAs at a global level in oocytes.

Nanog is known for its prominent function as a regulator of pluripotency in embryonic stem cells (Mitsui et al., 2003; Boyer et al., 2005; Loh et al., 2006) and reprogramming of somatic cells to the pluripotent state (Takahashi and Yamanaka, 2006; Silva et al., 2009). In zebrafish, Nanog has been shown to be a transcriptional activator to play a central role in regulating early embryogenesis. For instance, maternal Nanog mediates endoderm formation through the Mxtx2-Nodal signaling (Xu et al., 2012), and is required for both extra-embryonic development (Gagnon et al., 2017) and embryonic architecture formation (Veil et al., 2018). During the maternal-to-zygotic transition (MZT), the maternally provided transcription factors, Pou5f3, SoxB1, and Nanog coordinately open up chromatin to initiate zygotic genes activation (Lee et al., 2013; Veil et al., 2019; Palfy et al., 2020). Our recent study shows that Nanog represses the global activation of maternal β-catenin activity to safeguard the dorsal-ventral axis formation (He et al., 2020). However, as a strongly maternally expressed gene, the role of Nanog in oocyte development and maturation is still unknown.

In this study, we found that the absence of maternal *nanog* led to various developmental defects in oocytes and early embryos. Our study demonstrates the global translational activity is greatly enhanced in *nanog* mutant oocytes and maternal *nanog* mutant (M*nanog*) embryos, due to the transcriptional activation of *eukaryotic translation elongation factor 1 alpha 1, like 2* (*eef1a1l2*) during oocyte maturation. We further show that maternal depletion of *eef1a1l2* significantly rescues the developmental defects of *nanog* mutant oocytes and early development of M*nanog* embryos. Thus, our study reveals a novel role for zebrafish Nanog as a general translational repressor through transcriptional repression of *eef1a1l2* in oocyte maturation.

## Results

### Maternal *nanog* is required for oocyte maturation and early embryonic development

We have obtained *nanog* mutant using TALEN technology and addressed its critical role in regulating dorsal formation through interfering with TCF factors in previous studies (He et al., 2015; He et al., 2020). Through analyzing the phenotype of M*nanog* embryos, we found that the M*nanog* embryos showed slow epibolic movement, resulting in the accumulation of blastomere cells at the animal pole at gastrulation stage (Fig. 1A), which is similar to the phenotype of maternal and zygotic mutant of *nanog* (MZ*nanog*) (He et al., 2020). Comparing the size of M*nanog* and WT embryos at 15 minutes post fertilization (mpf), the M*nanog* embryos showed significantly shorter chorion diameter and oocyte diameter than WT embryos (Fig. 1B-D). In addition, we analyzed the activation phenotype of mutant egg by monitoring the cortical granule (CG) exocytosis and cytoplasmic streaming. Fluorescein-conjugated *Maclura pomifera* lectin was used to label CGs to assess CG exocytosis in activated eggs. Compared with WT eggs, *nanog* mutant eggs showed abundantly retained CGs (Fig. 1E) at 10 minutes post activation (mpa). CellTracker™ CM-DiI Dye was injected into the yolk of M*nanog* and WT embryo to monitor cytoplasmic streaming. In WT embryos, vigorous cytoplasmic movement toward the animal pole was recorded (Movie 1). In contrast, M*nanog* embryos showed sluggish cytoplasmic streaming (Movie 2). These results indicate that maternal Nanog is essential for egg activation and early embryonic development.

**Fig. 1.**
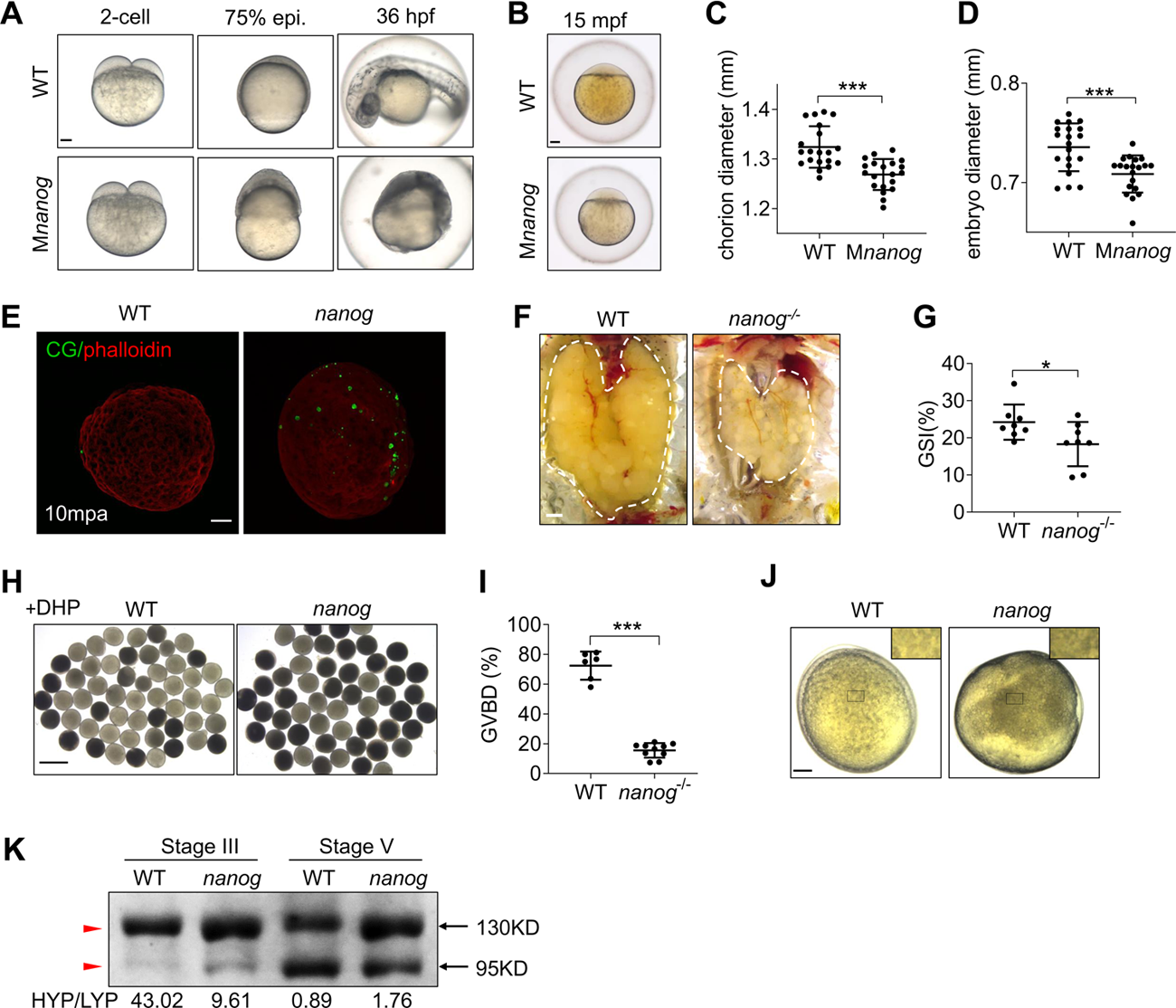
Loss of maternal *nanog* resulted in oocyte maturation defects. (A) Bright-field images showing the embryonic malformation of M*nanog* mutants in contrast to time-matched WT embryos. Scale bar, 100μm. (B) WT and M*nanog* embryos with chorions at 15 min post-fertilization (mpf). Scale bar, 100μm. (C, D) Measurement of chorion diameter and oocyte diameter at 15 mpf. \*\*\**P*<0.001. N=20. (E) Representative images showing labelling of cortical granules (CG) in WT and *nanog* mutant eggs fixed at 10 mpa. F-actin was stained using phalloidin to show the outline of embryo. Scale bar, 100μm. N=25. (F) Appearance of ovaries dissected from WT and *nanog*^-/-^ females. Scale bar, 1mm. (G) The GSI scatterplot of WT and *nanog*^-/-^ females. N=8. GSI, gonadosomatic index. \**P*<0.05. (H) Morphology of stage IV oocytes dissected from WT and *nanog*^-/-^ ovaries with incubation of 17α,20β-dihydroxy-4-pregnen-3-one (DHP, 1μg/mL) after 2 h. Scale bar, 1mm. (I) Comparison of the %GVBD in WT and *nanog* mutant oocytes. 6 WT and 10 *nanog*^-/-^ fishes were analyzed. (J) Stage V oocytes from WT and *nanog* mutant. Insets show enlarged regions of the yolk and relative opaqueness is seen in *nanog* mutants. Scale bar, 100μm. (K) SDS-PAGE and Coomassie staining of major yolk proteins of stage III oocytes and stage V oocytes. The higher and lower molecular weight yolk proteins (HYP and LYP) are indicated by the red arrowhead. HYP/LYP ratios were calculated to represent yolk protein cleavage levels.

Moreover, the TUNEL assay showed apoptotic signals appeared in Balbiani bodies and the cytoplasm of early-stage mutant oocytes, but not in WT oocytes (Fig. S1A). Mitochondria are enriched in the Balbiani body and also present throughout the oocyte cytoplasm (Marlow and Mullins, 2008; Jamieson-Lucy and Mullins, 2019), thus *nanog* deficiency induced the mitochondrial apoptosis during oocyte maturation. Besides, robust active-Caspase3 signals was also detected in M*nanog* embryos but not WT embryos at 75% epiboly stage (Fig. S1B). These data demonstrate *nanog* depletion induces the oocyte apoptosis and the death of early embryonic cells. Through morphological and histological analyses, we found the gonadosomatic index (GSI; gonad weight/body weight X 100%) was significantly reduced in *nanog*^-/-^ compared to WT females (Fig. 1F, G), indicating defects of oocyte maturation in *nanog* mutants. To furtherly characterize the defects of oocyte maturation of *nanog* mutant, stage IV oocytes were isolated and treated with 17α,20β-dihydroxy-4-pregnen-3-one (DHP) to determine the percentage of germinal vesicle breakdown (GVBD) *in vitro.* After 2 hours incubation, the percentage of GVBD in *nanog* mutant oocytes is 15.5%, which is significantly lower than that in WT oocytes (72.4%) (Fig. 1H, I). Moreover, the *nanog* mutant stage V oocyte showed less transparent than WT (Fig. 1J). During the oocyte maturation, the major yolk proteins undergo cleavage and change the appearance of oocyte from opaque to transparent (Dosch et al., 2004). So we compared the composition of the higher and lower molecular weight yolk proteins (HYP and LYP) from stage III to stage V oocytes, to evaluate the yolk protein cleavage level. In stage III oocytes, the HYP/LYP ratio were even lower in *nanog* mutant than in WT, whereas the ratio was greatly decreased in WT stage V oocytes than in mutants (Fig. 1K). These results suggested the deficiency of yolk protein cleavage during maturation of *nanog* mutant oocytes. In addition, we also generated a transgenic line of *Tg(CMV:nanog-myc*) in *nanog* homozygous background. Immunofluorescence staining showed Nanog is strongly expressed in early oocytes using anti-Myc antibody (Fig. S1C). Moreover, this overexpression of *nanog* could rescue the early developmental defects of M*nanog* (Fig. S1D), demonstrating that oocyte maturation defect and early embryonic defect were caused by deficiency of Nanog. Therefore, we conclude that loss of maternal *nanog* leads to pleiotropic defects of oocyte maturation and early embryonic development.

### Loss of maternal *nanog* elevates the global translation level

In order to understand the molecular mechanism of Nanog regulating oocyte maturation, we quantitatively compared proteomes of *nanog* mutant eggs with WT using isobaric tags for relative and absolute quantitation (iTRAQ) technology. More than 1600 proteins were identified in eggs from the two genotypes. The identified proteins were classified using COG database (Clusters of Orthologous Groups of proteins), and two top categories are involved in the protein translation-related biological process (cluster J and O) (Fig. S2A). Compared the mutant with WT, 67 proteins show differential expression (*P*<0.05) (S1Table), and 39 proteins were increased in mutant eggs (Fig. S2B). Gene ontology analysis of the upregulated proteins showed that one of the most significant enriched biological processes is translation elongation factor activity (Fig. S2C). Gene-Concept Network showed four elongation factors were enriched (Fig. S2D). These results imply that the global translation activity is elevated in *nanog* deficiency eggs.

To verify this speculation, we assessed the translation activity of M*nanog* embryos at early developmental stage. The mCherry reporter mRNA was injected into one-cell stage WT and M*nanog* embryos together with the same amount of GFP protein. The injected embryos were imaged under a fluorescence microscope at later stages and fluorescence levels were measured. The GFP protein acted as the loading control for injection. Fluorescence measurement showed that the reporter mRNA translation level was significantly higher in M*nanog* embryos (Fig. 2A, B). Using western blotting analysis, we confirmed the increase of mCherry reporter translation in maternal *nanog*-depleted embryos (Fig. 2C, D). More interesting is, we treated the stage IV mutant oocytes with an inhibitor of translation initiation factor, *eif4a*, and determined the percentage of GVBD in vitro. After 2 hours incubation, the percentage of GVBD was remarkably increased from 10.4% to 28.3% in mutant oocytes (Fig. S2E, F). Taken together, these results suggest that Nanog is necessary for repressing the global translation of maternal mRNAs during oocyte maturation and early embryonic development.

**Fig. 2.**
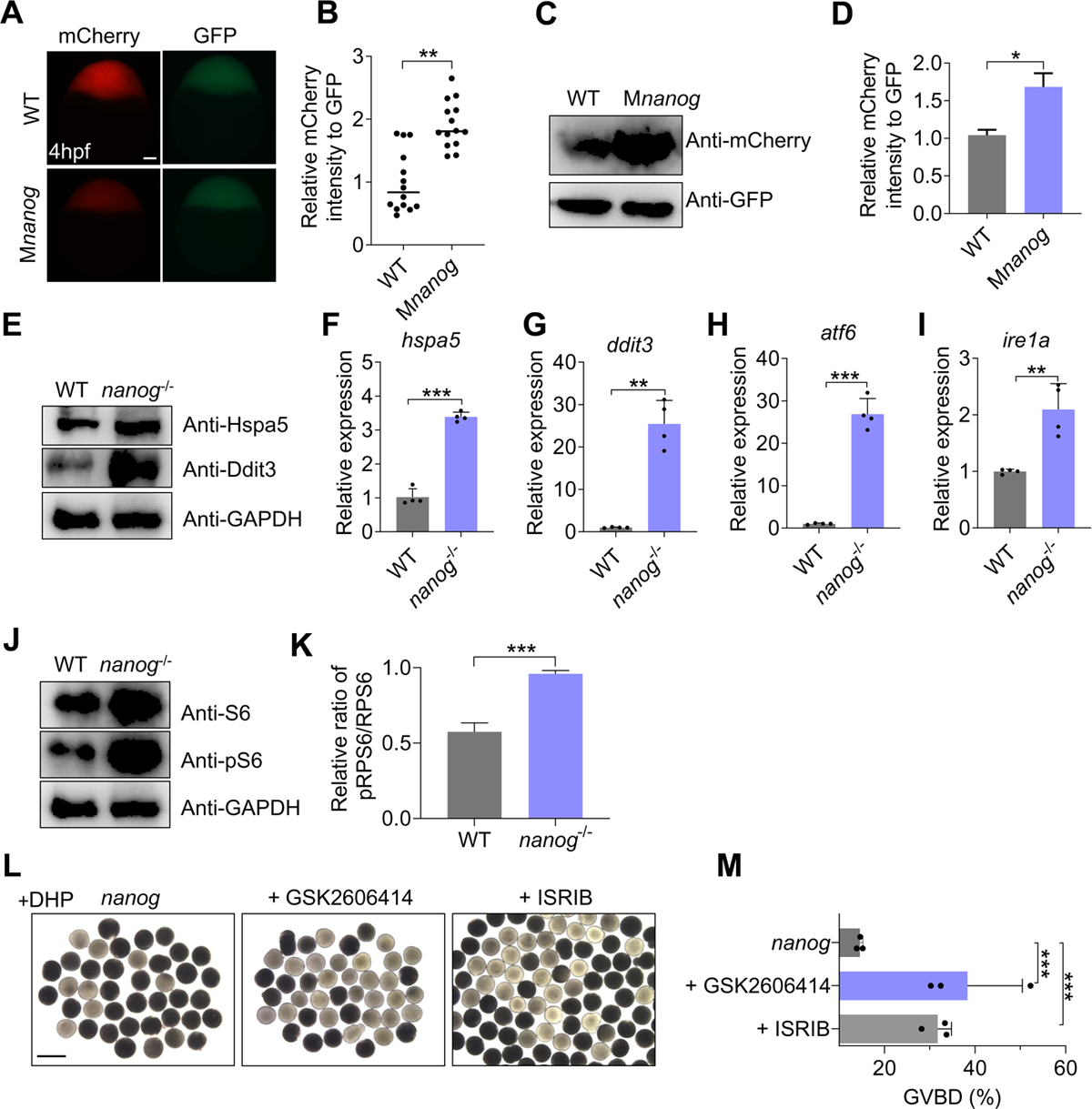
Loss of *nanog* triggers ER stress/unfolded protein response (UPR) and elevates global translation activity. (A) Fluorescent images showing mCherry reporter levels with GFP protein control levels in WT and M*nanog* embryos at 4hpf. Scale bar, 100μm. (B) Measurement of mCherry reporter intensities relative to GFP. \*\**P*<0.01. n=14. (C, D) Western blotting analysis of mCherry reporter levels at 4 hpf. \**P*<0.05; Student’s t-test. (E) Western blot analysis showed increased expression of Hspa5 (Bip) and Ddit3 (CHOP) in *nanog*^-/-^ ovary. (F-I) RT-qPCR analysis of *hsp5a* (F), *ddit3* (G), *atf6* (H) and *ire1a* (I) in WT and *nanog*^-/-^ ovaries. \*\**P*<0.01, \*\*\**P*<0.001. (J) Western blot analysis of S6 and phosphorylated S6 in WT and *nanog*^-/-^ ovary. (K) statistical analysis of phosphorylated S6/S6 level in panel J. (L) Morphology of stage IV oocytes dissected from *nanog*^-/-^ ovaries and treated with PERK inhibitor after 2h. Oocytes dissected from 3 fishes were treated for 2 hours in the presence of DHP. Final concentration: GSK2606414, 50 nM; ISRIB, 5 μM; DHP: 1μg/mL. Scale bar, 1mm. (M) %GVBD comparison in *nanog* mutant oocytes treated with or without PERK inhibitor. n=3. \*\**P*<0.01, \*\*\**P*<0.001.

### Nanog depletion triggers ER stress and the unfolded protein response (UPR)

The endoplasmic reticulum (ER) functions as a crucial machinery for protein synthesis, modification and trafficking in eukaryotic cells. Under ER stress, cells activate the UPR to alleviate ER burden by reducing protein translation, increasing protein degradation and generating additional chaperones to assist protein folding. Therefore, ER stress and the UPR are often associated with aberrant translational derepression (Kaufman, 2002; Miao et al., 2017). The UPR functions through three major pathways, initiated by three ER-localized transmembrane proteins, such as protein kinase RNA-like ER kinase (PERK), activating transcription factor 6 (ATF6), and Inositol-requiring enzyme 1 (IRE1), to maintain ER homeostasis (Hetz, 2012). Normally, the N-terminal of these transmembrane ER proteins are held by ER chaperone Hspa5 (also termed Grp78 or Bip), preventing their aggregation. When misfolded proteins accumulate, Hspa5 releases, allowing aggregation of these transmembrane signaling proteins, and launching the UPR (Rao and Bredesen, 2004; Shen et al., 2004; Schroder and Kaufman, 2005). Activation of PERK upregulates the expression of CCAAT-enhancer-binding protein homologous protein (CHOP), which induces cell apoptosis and death (Oyadomari and Mori, 2004; Iurlaro and Muñoz-Pinedo, 2016). We detected the transcriptional level and protein level of *hspa5* and CHOP-encoding gene *ddit3* in mutant ovary, found that both transcription and translation of *hspa5* and *ddit3* are increased in *nanog*^-/-^ ovary (Fig. 2, E-G). TUNEL assay also showed obvious apoptosis signal in mutant ovary (Fig. S1A). Due to lack of specific antibodies against zebrafish ATF6 and IRE1A, we detected the mRNA expression of *atf6* and *ire1a*, and discovered that both of *atf6* and *ire1a* expression were increased in mutant oocytes (Fig. 2H, I). Since phosphorylated ribosomal protein S6 (pS6) is considered an indicator of active protein synthesis (Biever et al., 2015; Meyuhas, 2015), we also detected the expression level of total S6 and pS6, and found that the pS6 level was significantly increased in *nanog* mutant ovary (Fig. 2J, K). Finally, we used PERK inhibitors (GSK2606414 and ISRIB) to treat *nanog* mutant oocytes and determined the occurrence of GVBD. In both treatments, the percentage of GVBD of mutant oocytes were recovered (Fig. 2L, M). These results demonstrate that loss of *nanog* triggers ER stress and UPR and thus leads to failure of oocyte maturation.

### Loss of maternal Nanog up-regulates *eef1a1l2* transcription level

To find out the molecular mechanism responsible for Nanog regulating translation activity in early embryos, we conducted RNA sequencing (RNA-seq) analysis between *nanog* mutant eggs and WT eggs (accession number: PRJNA633216). A total of 207 genes are differentially expressed between mutant and WT eggs, with 102 upregulated genes and 105 downregulated genes in mutant eggs. Among the upregulated genes, the difference of *eukaryotic elongation factor 1 alpha 1, like 2* (*eef1a1l2*) is among the most significant ones (Fig. 3A). The eEF1A1l2 is a major subunit of the translation elongation factor 1 complex (eEF1), which plays a central role in protein synthesis by delivering aminoacyl-tRNAs to the elongating ribosome (Sasikumar et al., 2012). RPKM calculation and visualization of RNA-seq reads mapped to the *eef1a1l2* showed significant upregulation of *eef1a1l2* in mutant eggs (Fig. S3A, B). Reverse-transcription quantitative PCR (RT-qPCR) further proved that the transcription of *eef1a1l2* was significantly increased in *nanog* mutant oocytes at five different stages and in early embryonic developmental stages (Fig. 3B, S3C). In situ hybridization of ovary cryosection also showed increased expression of *eef1a1l2* in *nanog* mutant (Fig. 3C). In contrast, transgenic overexpression of *nanog* significantly decreased the high expression levels of *eef1a1l2* in mutant oocytes (Fig. 3B, C). These data indicate that *nanog* deficiency led to strong transcriptional activation of *eef1a1l2* during oocyte maturation.

**Fig. 3.**
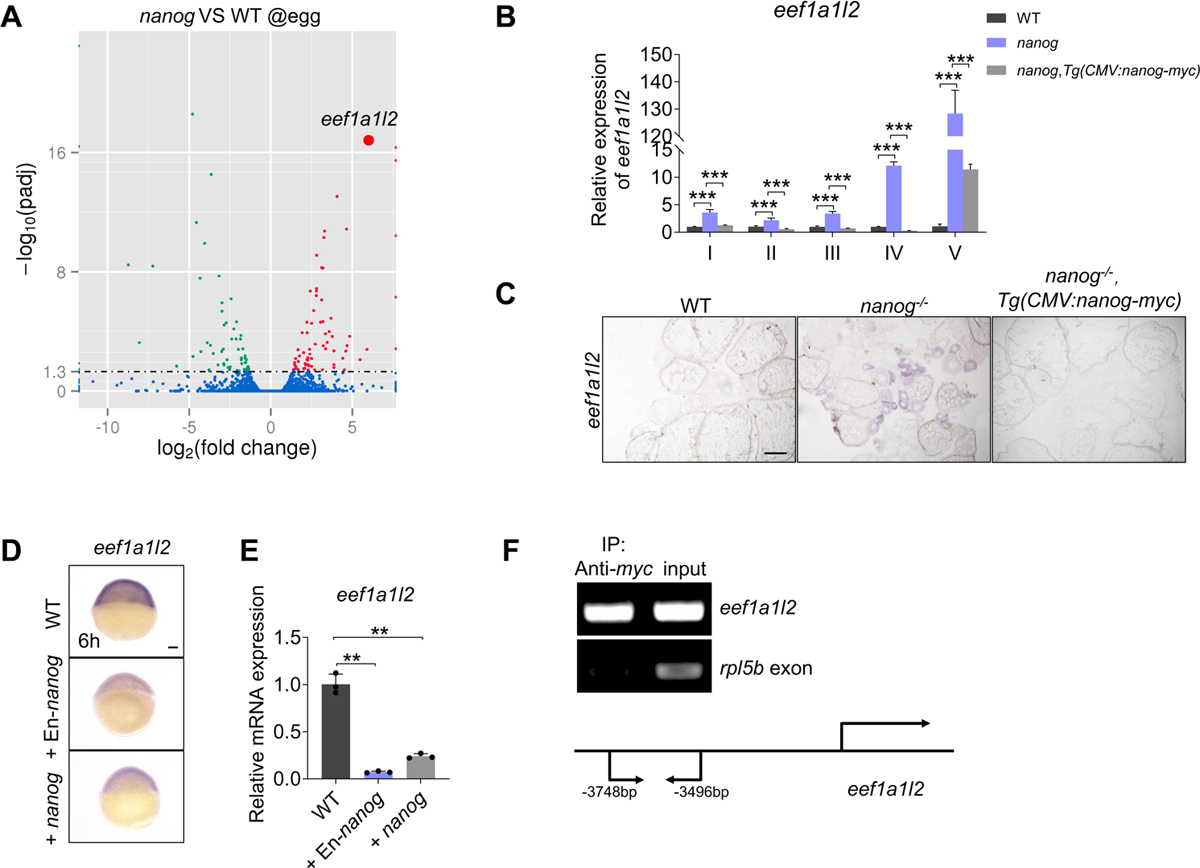
Nanog transcriptionally inhibits the expression of *eef1a1l2*. (A) RNA-seq analysis showed significantly increased expression of *eef1a1l2* in *nanog* null egg. Red dots indicate upregulated genes, green dots indicate down-regulated genes, and blue dots indicate no difference genes in *nanog* mutant egg. (B) Upregulated expression of *eef1a1l2* in *nanog* mutant oocytes at different stages revealed by RT-qPCR. \*\**P*<0.01, \*\*\**P*<0.001. (C) In situ hybridization on cryosections of ovaries and WISH analysis of unfertilized eggs revealed increased expression of *eef1a1l2* in *nanog* mutant. Scale bar, 200μm. (D,E) WISH (D) and RT-qPCR (E) analysis showed reduced expression of *eef1a1l2* in *nanog* or En-*nanog* overexpressed embryos at 6 hpf. \*\**P*<0.01. Scale bar, 100μm. (F) ChIP analysis of *Tg*(CMV:*nanog-myc*) ovaries with anti-Myc antibody at 6mpf. The promoter region of *eef1a1l2* was enriched in precipitated chromatin. *rpl5*b was served as a negative control.

To verify the transcriptional inhibition of *eef1a1l2* by Nanog, wildtype *nanog* or a constitutive repressor type *nanog* (Engrailed fusion with Nanog homeodomain, En-*nanog*) (He et al., 2020) was overexpressed and the transcription of *eef1a1l2* was measured at shield stage. Both in situ hybridization and RT-qPCR analysis showed that the expression of *eef1a1l2* was significantly reduced in both *nanog* and En-*nanog* overexpressed-embryos (Fig. 3D, E), suggesting that Nanog acts as a transcriptional repressor on the regulation of *eef1al2*. To clarify whether Nanog directly bind to the promoter region of *eef1a1l2* to repress its transcription, a chromatin immunoprecipitation (ChIP) assay was conducted. Ovaries of *Tg*(CMV:*nanog-myc*) at 6mpf were dissected and ChIP was performed using anti-Myc antibody. The precipitated chromatin was then analyzed by PCR using primer pairs that could amplify fragments of *eef1a1l2* promoter. A fragment of the *rpl5b* exon amplified by a specific primer pair was used as control (Belting et al., 2011). As shown in Fig. 3F, promoter fragment of *eef1a1l2* was significantly enriched in the immunoprecipitated while no enrichment of the control genomic region *rpl5*. Thus, this result demonstrates that Nanog directly binds to the promoter of *eef1a1l2* to inhibit its transcription. All these data illustrate that Nanog directly inhibits the transcription of *eef1a1l2* and depletion of *nanog* leads to significantly increased expression of *eef1a1l2*.

### Deficiency of eEF1A1l2 ameliorates impaired oocyte maturation of *nanog* mutants

We then generated homozygous mutant of *eef1a1l2* and two types of *eef1a1l2* mutants were obtained and neither of them showed obvious defect (Fig. S4), and we used the ihb99 allele for subsequent study. To determine if Nanog promotes oocyte maturation through suppression of *eef1a1l2*, we generated double homozygous mutant of *nanog* and *eef1a1l2* and studied whether depletion of *eef1a1l2* could ameliorate the oocyte maturation defect of *nanog* mutant. Through morphological analysis, we found double maternal mutant (M*nanog*, M*eef1a1l2*) showed increased embryo chorion diameter and oocyte diameter at 15 mpf, compared to M*nanog* embryos (Fig. 4A-C). The process of cortical granule (CG) exocytosis in double mutant eggs (*nanog*,*eef1a1l2*) were also comparable to WT at 10 mpa, which showed less retained CGs than *nanog* mutant (Fig. 4D). Moreover, cytoplasmic streaming labeled by CM-DiI Dye in double maternal mutant embryo (M*nanog*, M*eef1a1l2*) showed vigorous cytoplasmic movement, similar to the WT (Movie 3). These results indicate that depletion of *eef1a1l2* rescues the egg activation defect of *nanog* mutant.

**Fig. 4.**
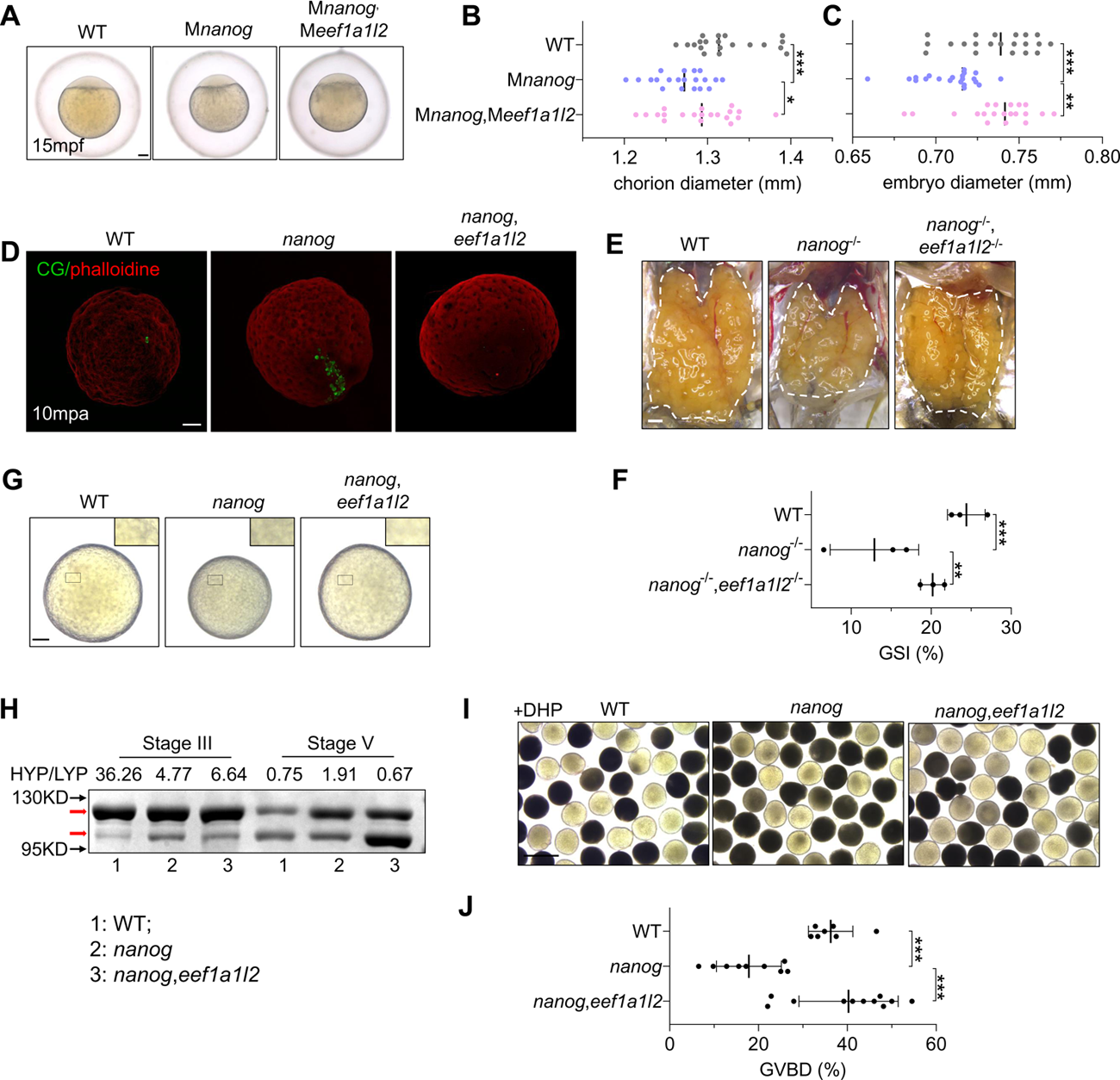
Depletion of *eef1a1l2* rescues the impaired oocyte maturation of *nanog* mutant. (A) WT, M*nanog* and M*nanog*, M*eef1a1l2* embryos with chorions at 15 min post-fertilization (mpf). Scale bar, 100μm. (B,C) Measurement of chorion diameters and oocyte diameters at 15 mpf. \**P*<0.05, \*\**P*<0.01, \*\*\**P*<0.001. n=20. (D) Representative images showing labelling of cortical granules (CG) in WT, *nanog*, and *nanog*-*eef1a1l2* double mutant eggs fixed at 10 mpa. F-actin was stained using phalloidin to show the outline of embryo. n=15. Scale bar, 100μm. (E) Appearance of ovaries dissected from WT, *nanog*^-/-^ and *nanog*^-/-^,*eef1a1l2*^-/-^ females. Scale bar, 1mm. (F) The GSI scatterplot of WT, *nanog*^-/-^ and *nanog*^-/-^,*eef1a1l2*^-/-^ females. n=3. GSI, gonadosomatic index. \*\**P*<0.01, \*\*\**P*<0.001. (G) Morphology of eggs from WT, *nanog*^-/-^ and *nanog*^-/-^,*eef1a1l2*^-/-^. Insets show enlarged regions of the yolk. Scale bar, 100μm. (H) SDS-PAGE and Coomassie staining of major yolk proteins of stage III oocytes and stage V eggs. The higher and lower molecular weight yolk proteins (HYP and LYP) are indicated by the arrow. HYP/LYP ratios were calculated to represent yolk protein cleavage levels. (I) Morphology of stage IV oocytes dissected from WT, *nanog*^-/-^ and *nanog*^-/-^,*eef1a1l2*^-/-^ ovaries with incubation of DHP (1μg/mL) after 2h. Scale bar, 1mm. (J) Comparison of the %GVBD in WT, *nanog*^-/-^ and *nanog*^-/-^,*eef1a1l2*^-/-^ oocytes. WT n=11, *nanog*^-/-^ n=9, *nanog*^-/-^,*eef1a1l2*^-/-^ n=7.

Therefore, we further examined the oocyte maturation improvement in double mutant. Morphologically, The GSI was increased in double mutant (*nanog*^-/-^, *eef1a1l2*^-/-^) female, compared to *nanog*^-/-^ (Fig. 4E, F). The double mutant eggs showed more transparent than *nanog* mutant, similar to WT (Fig. 4G). The altered ratio of the higher and lower molecular weight yolk proteins (HYP and LYP) from stage III oocytes to stage V oocytes in double mutant confirmed this conclusion (Fig. 4H). The % GVBD was also remarkably increased in double mutant oocytes (Fig. 4I, J). These results together indicate that Nanog promotes oocyte maturation through suppressing the expression of *eef1a1l2* in oocyte.

### Depletion of *eef1a1l2* alleviates the ER stress and UPR in *nanog* mutant oocyte

Since depletion of *eef1a1l2* could ameliorates the oocyte maturation defect of *nanog* mutant, we wondered the eEF1A1l2 depletion on alleviating of ER stress. We addressed the mRNA expression level of ER stress-associated genes, found that the increased expression of *hspa5*, *ddit3*, *atf6* and *ire1a* in *nanog* mutant ovary were all restored in *nanog* and *eef1a1l2* double mutant (Fig. 5A-D). IRE1 is a unique RNase which removes an internal 26 nucleotides from X-box binding protein 1 (XBP1) mRNA transcripts in the cytoplasm and activates gene expression involved in protein folding and degradation, thus the splicing of XBP1 mRNA has been established as a common indicator of ER stress (Shen et al., 2001; Yoshida et al., 2001; Li et al., 2015). We detected the altered splicing of *xbp1* in ovary of mutant and WT, discovered the splicing ratio of *xbp1* was increased in *nanog* mutant ovary, and this excessive splicing was reduced in double mutant ovary (Fig. 5E, F).

**Fig. 5.**
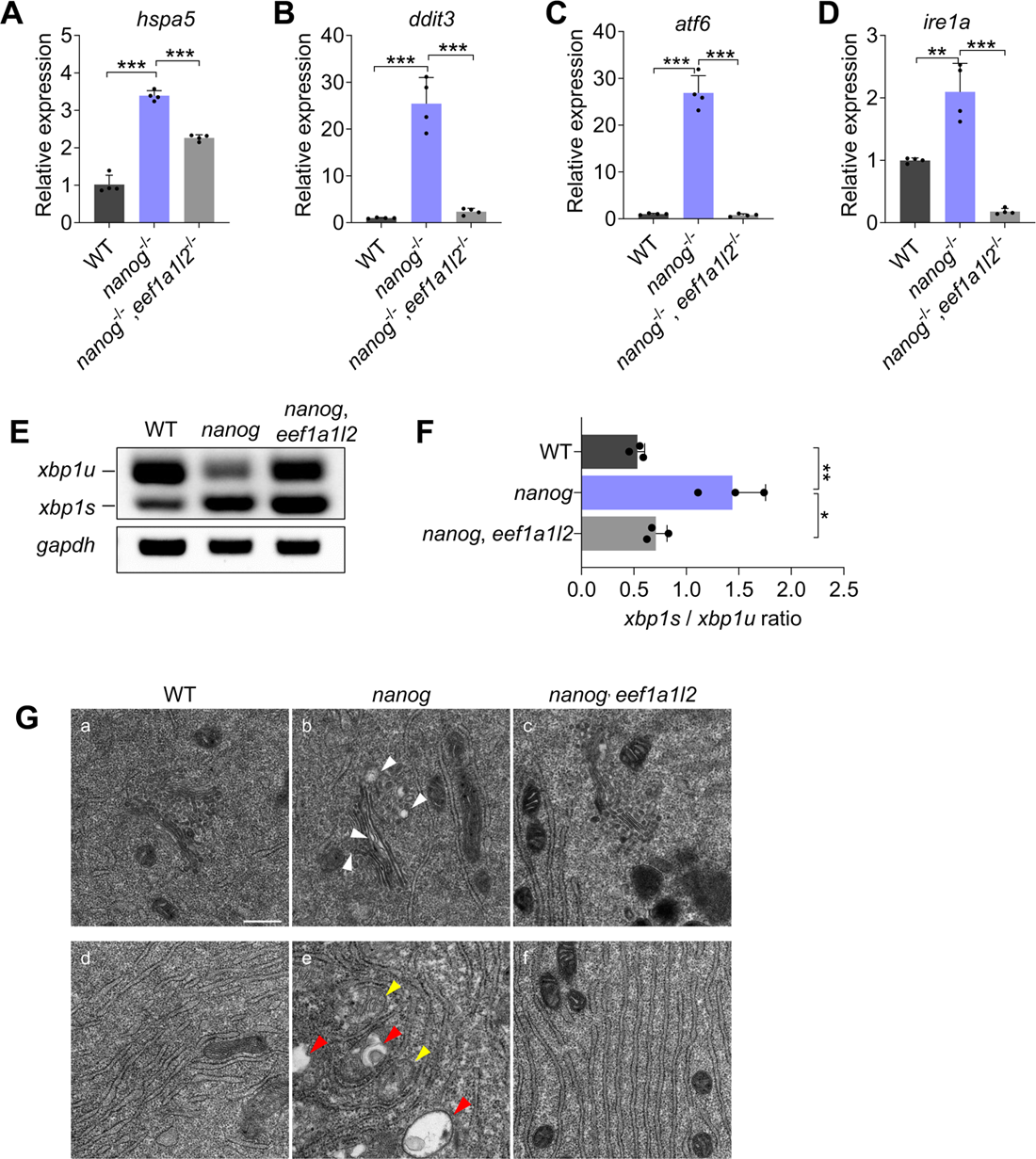
Depletion of *eef1a1l2* ameliorates ER stress and UPR of *nanog* mutant oocyte. (A-D) RT-qPCR analysis showed expression reduction of *hspa5* (A), *ddit3* (B), *atf6* (C) and *ire1a* (D) in *nanog* and *eef1a1l2* double mutant ovary, compared with *nanog* mutant ovary. \**P*<0.05, \*\**P*<0.01, \*\*\**P*<0.001. (E, F) RT-PCR examination of *xbp1* splicing. The ratio of spliced *xbp1* (*xbp1s*) mRNA to unspliced *xbp1* (*xbp1u*) mRNA was increased in *nanog* mutant ovary, but restored in *nanog* and *eef1a1l2* double mutant. *gapdh* is used as internal control. The *xbp1s*/*xbp1u* ratio in F represents the intensity ratio of corresponding PCR product bands in E. (G) Observation of ER, Golgi and mitochondria structure in WT, *nanog* mutant and double mutant at stage I oocytes by transmission electron microscopy. White arrowheads indicate Golgi apparatus, yellow arrowheads indicate mitochondria, red arrowheads indicate lysosome. Scale bar, 0.5 μm.

To observe organelle changes that may accompany response to ER stress, ultrastructure analysis using transmission electron microscopy (TEM) was performed in stage I oocytes of WT, *nanog* mutant, and double mutant. The WT oocytes showed normal morphology of ER, Golgi apparatus and mitochondria (Fig. 5G, a and d). However, *nanog* mutant oocytes showed disruption of Golgi apparatus, including swelling of Golgi apparatus, dilated and disintegrated vesicles, and collapse of Golgi complex (Fig. 5G, b, white arrowheads). *nanog* mutant oocytes also showed incompact and swollen mitochondria (Fig. 5G, e, yellow arrowheads), as well as evident lysosome distribution (Figure 5G, e, red arrowheads). In contrast, *nanog* and *eef1a1l2* double mutant showed well structure of ER, Golgi apparatus and mitochondria (Figure 5G, c, f). These data demonstrate that depletion of *eef1a1l2* alleviates the ER stress and UPR in *nanog* mutant, indicating that transcriptional activation of *eef1a1l2* in *nanog* mutant oocytes inducing ER stress and UPR, thus leading to defects of oocyte maturation.

### Depletion of *eef1a1l2* rescues early embryonic development defect of *nanog* mutant

Given that depletion of *eef1a1l2* could rescue the oocyte maturation defect and alleviate the ER stress/UPR, we wondered the rescue effect of depletion of *eef1a1l2* on early embryonic development defect. Due to the translation elongation role of eEF1A1l2, we firstly assessed the translation level of double mutant. mCherry mRNA reporter and GFP protein were co-injected at one-cell stage in WT, M*nanog*, and M*nanog*,M*eef1a1l2* embryos. Fluorescence intensity of mCherry was measured at 4 hpf. Fluorescence measurement showed that the reporter translation level was significantly reduced in M*nanog*,M*eef1a1l2* embryos (Fig. S5A, B), suggesting the elevated translation level of *nanog* mutant was decreased by depletion of *eef1a1l2*.

Next, we examined the early embryonic development phenotype of double mutant. The morphological phenotypes of WT, M*nanog* (*nanog*^-/-^ female cross with WT male), M*nanog*,M*eef1a1l2* (*nanog*^-/-^,*eef1a1l2*^-/-^ female cross with WT male), M*nanog*,MZ*eef1a1l2* (*nanog*^-/-^,*eef1a1l2*^-/-^ female cross with *eef1a1l2*^-/-^ male), and M*nanog*,Z*eef1a1l2* (*nanog*^-/-^,*eef1a1l2*^+/-^ female cross with *eef1a1l2*^-/-^ male and *nanog*^+/-^,*eef1a1l2*^-/-^ embryos were genotyped) were recorded at 0.2, 8, 12 and 24 hpf, respectively. As described in Fig. 1A, blastomere cells stacked at the animal pole and unable to complete the gastrulation in M*nanog* embryos. Until 24 hpf, M*nanog* embryos still showed abnormal shapes and died gradually (Fig. 6A). Surprisingly, either getting rid of maternal increased *eef1a1l2* in M*nanog*,M*eef1a1l2* embryos, or eliminating both maternal and zygotic increased *eef1a1l2* in M*nanog*,MZ*eef1a1l2* embryos, could effectively rescue the developmental defects of M*nanog* (Fig. 6A). M*nanog*,M*eef1a1l2* and M*nanog*,MZ*eef1a1l2* embryos both exhibited ameliorated epiboly movement at gastrulation stage and improved axial formation excepting telencephalon defect at prim-stage. However, M*nanog*,Z*eef1a1l2* embryos only lacking the zygotic *eef1a1l2* exhibited similar phenotype with M*nanog* (Fig. 6A), indicating that the maternally provided *eef1a1l2* mRNA in M*nanog*,Z*eef1a1l2* still led to developmental failure. In summary, only if *eef1a1l2* was completely disrupted in the *nanog* mutant oocytes, the M*nanog*,M*eef1a1l2* or M*nanog*,MZ*eef1a1l2* embryos displayed rescued early embryonic development. These data indicate that support the c maternal activation of *eef1a1l2* in *nanog* mutant oocytes not only leads to oocyte maturation defects, but also results in early developmental defects of M*nanog* embryos.

**Fig. 6.**
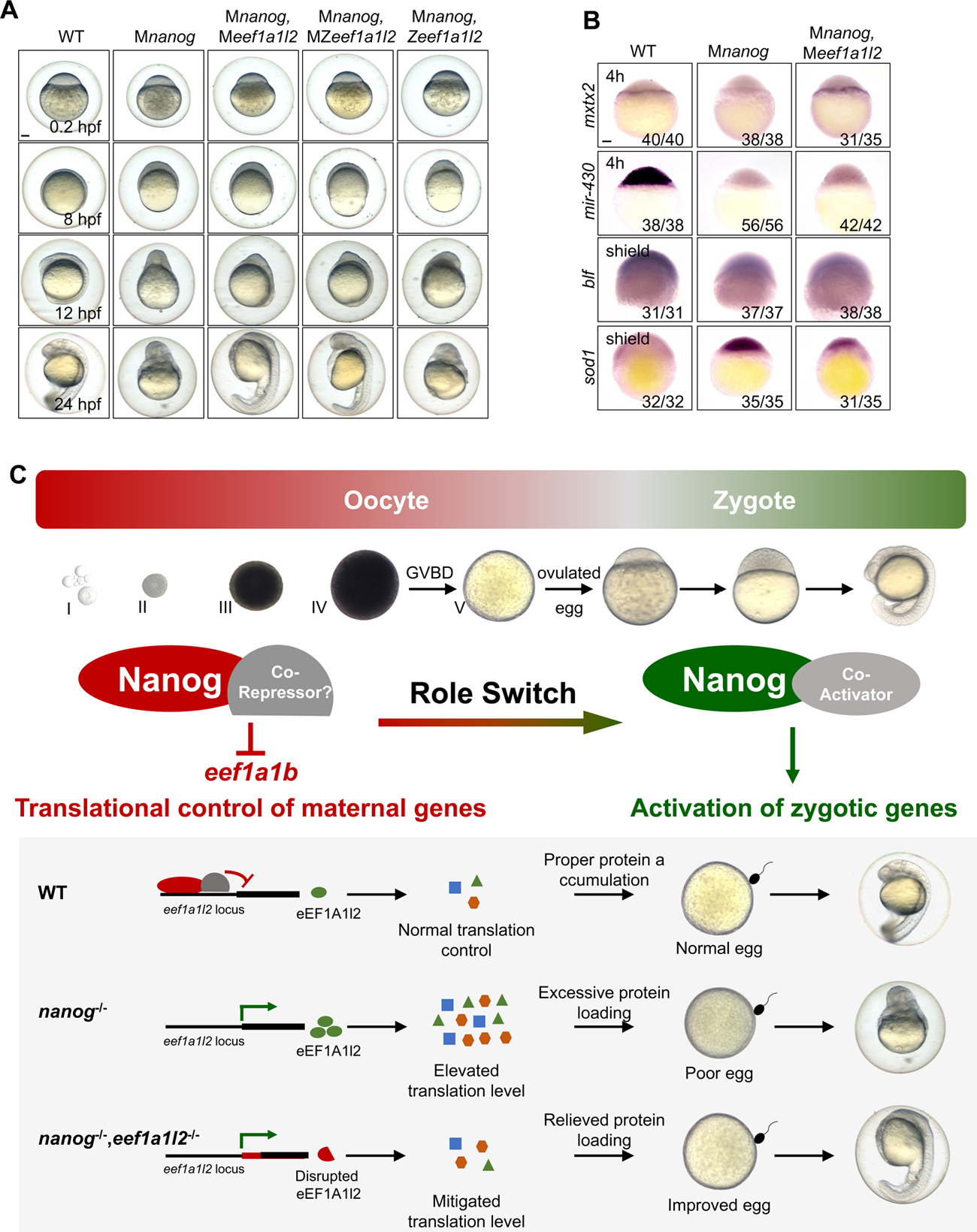
Maternal depletion of *eef1a1l2* rescues early embryonic defects of *nanog* mutant. (A) Phenotype of WT, M*nanog*, M*nanog*,M*eef1a1l2*, M*nanog*,MZ*eef1a1l2* and M*nanog*,Z*eef1a1l2,* embryos at 0.2, 8, 12, and 24 hpf. Scale bar, 100 μm. (B) Detection of *mxtx2*, *mir-430* precursor, *blf* and *sod1* in WT, M*nanog* and M*nanog*,M*eef1a1l2* embryos by WISH at indicated stage. scale bar, 100 μm. (C) The model of Nanog function in oocyte and early embryo of zebrafish. Top: Nanog acts as a transcriptional repressor to suppress the expression of *eef1a1l2* and maintain the correct translation level of maternal mRNAs during oocyte development. Then Nanog switches to a transcriptional activator to prime zygotic genome activation in zygotes. Bottom: in WT oocyte, Nanog inhibits the transcription of *eef1a1l2* and maintain the proper level of global translation, ensuring appropriate amounts of proteins. Good egg quality and normal embryonic development is thus guaranteed. In the absence of maternal *nanog* (*nanog*^-/-^), the balance of global translation is destroyed. Elevated translation resulted in excessive protein loading, further leads to poor egg quality and failure embryogenesis. In *nanog* and *eef1a1l2* double mutant oocyte (*nanog*^-/-,^ *eef1a1l2*^-/-^), global translation level is mitigated due to the absent of eEF1A1l2, protein overloading is relieved and egg quality is also improved, thereby promotes a better embryonic morphology formation.

Furthermore, we investigated the rescue effects using a set of molecular markers representing different functions of Nanog in early development. Several studies have proved that maternal Nanog directly activates *mxtx2* to regulate endoderm and extraembryonic formation through Nanog-*mxtx2*-Nodal pathway (Xu et al., 2012; Gagnon et al., 2018; Veil et al., 2018).We confirmed that the expression of *mxtx2* was absent in M*nanog*, while the disappearance of *mxtx2* transcripts could be restored in M*nanog*,M*eef1a1l2* (Fig. 6B, S5C). During zebrafish zygotic genome activation (ZGA), together with Pou5f3 and SoxB1, maternal Nanog initiates the transcription of the first major wave of zygotic genes and directly activates microRNA miR-430, maternal mRNAs were then cleared by miR-430 at the post-ZGA (Giraldez et al., 2006; Lee et al., 2013). The expression of zygotic genes, miR-430 and *blf*, failed to be activated, and maternal mRNA *sod1*, which is targeted by miR-430, also failed to be removed at post-ZGA in M*nanog* (Fig. 6B, S5D-F). In contrast to rescue of *mxtx2* expression, the defects of ZGA and maternal mRNA clearance could not be rescued in M*nanog*,M*eef1a1l2* embryos (Fig. 6B, S5D-F). These results illustrate that during oogenesis, maternal Nanog safeguards the oocyte maturation by suppression the transcriptional activation of *eef1a1l2* as a transcriptional repressor, and during early embryogenesis, the transcriptional suppression of *eef1a1l2* by Nanog is mainly required for transcription initiation of *mxtx2* and YSL formation.

All these results deciphered a following molecular picture: during oocyte maturation and early embryogenesis (Fig. 6C). During WT oogenesis, Nanog acts as a transcriptional repressor, with certain co-repressors, directly inhibits the transcription of eukaryotic translation elongation factor, *eef1a1l2*, contributing to the translational control in oocytes. After fertilization, Nanog acts as a transcriptional activator, together with Pou5f3 and SoxB1, to initiate zygotic genome activation. In the oocytes produced by *nanog*^-/-^ females, the transcriptional inhibition of *eef1a1l2* is absent, ectopic maternal proteins are translated and accumulated, thus inducing ER stress and excessive UPR, and leading to oocyte maturation defects. The embryos fertilized from the *nanog*^-/-^ defective eggs failed to undergo normal gastrulation. In the oocytes produced by *nanog*^-/-^,*eef1a1l2*^-/-^ females, however, the translation level of maternal mRNAs is mitigated owing to the lack of functional eEF1A1l2, and ER stress and UPR are alleviated, therefore leading to a normal oocyte maturation. Therefore, it is likely that Nanog shifts from a transcriptional repressor to a transcription activator during oocyte-to-zygote transition (OZT), which are both essential for early embryogenesis.

## Discussion

The function of Nanog at early embryonic developmental stages has been well characterized in previous studies(Veil et al., 2018; Veil et al., 2019; He et al., 2020; Palfy et al., 2020). However, as a maternal expressed gene, its role in oocyte development and maturation is still unknown. In this study, we revealed that zebrafish Nanog is essential for oocyte maturation, which function has a lasting effect on early embryogenesis. Loss of maternal Nanog causes impaired oocyte maturation, deficient egg activation, and early embryo developmental failure. Impaired oocyte maturation correspondingly tends to produce poor-quality egg. We have compared the *nanog* expression level in good-quality egg and poor-quality egg and found that *nanog* expression is significantly decreased in poor-quality egg (Fig. S6), suggesting that Nanog might be a new factor for oocyte quality assessment. Mechanistically, Nanog transcriptionally represses the expression of translation elongation factor *eef1a1l2* to maintain a translational control state during oocyte maturation. In contrast, in *nanog* mutant oocytes, ectopic transcriptional activation of *eef1a1l2* elevates global translation activity and causes ER stress and UPR. Depletion of *eef1a1l2* could rescue the oocyte maturation defects and embryonic defects of *nanog* mutant. Taken together, our results delineate the mechanisms underlying a general translation repressor role of Nanog during oocyte maturation.

Maternal mRNAs synthesized in oocyte initiate the development of future generations. Some maternal mRNAs are either somatic or germline determinants and must be translationally repressed until embryogenesis (Richter and Lasko, 2011; Flora et al., 2018). Previous studies have provided numerous examples of how sequence-specific regulators, mostly RNA-binding proteins, maintain translational repression levels of mRNAs containing targeted cis-regulatory elements post-transcriptionally in gametogenesis and early embryogenesis (Kugler and Lasko, 2009; Kotani et al., 2013; Winata et al., 2017; Sun et al., 2018). In this study, we showed a transcriptional factor, Nanog, transcriptionally inhibits the expression of a translation elongation factor, eEF1A1l2 consequently controls the global mRNA translation activity during oocyte maturation. This translation repressive manner is realized through regulating the activity of a translation machinery, which is more global and independent on the sequence specificity of mRNA. Thus, this study revealed a novel translational control mechanism regulated by Nanog to promote oocyte maturation and early embryogenesis.

The genes who are not transcribed in oocyte and maturate eggs are considered as non-maternal genes, or zygotic genes. In theory, the transcription of non-maternal genes should be suppressed in oocyte to safeguard the oocyte maturation and early embryo patterning. The abnormal activation and expression of non-maternal genes will change the oocyte cell fate, affect egg quality, even lead to the apoptosis of oocyte. According to the expression pattern of *eef1a1l2* addressed during oocyte maturation and early embryonic development in WT (Fig. 3, S4), we found that *eef1a1l2* has no maternal expression in oocytes, indicating *eef1a1l2* is a non-maternal gene and not employed during oocyte maturation. However, *eef1a1l2* is precociously transcribed in *nanog* mutant oocytes, which leads to the over-activation of global translation activity, further results in the impaired oocyte maturation. This finding implies us Nanog safeguards oocyte maturation and early embryogenesis possibly by suppression the transcriptional activation of non-maternal genes.

Studies in human and mouse embryonic stem cells (ESC) revealed Nanog acts as both transcriptional activator and transcriptional repressor (Boyer et al., 2005; Liang et al., 2008). As a transcriptional activator, Nanog cooperating with Pou5f3 and Sox2, transcriptionally activates the expression of genes responsible for stem cell self-renew and pluripotency maintenance, while as a transcriptional repressor, Nanog associates with repression complexes and transcriptionally represses the expression of genes related to differentiation and development. In this study, we concluded that Nanog acts as a transcriptional repressor to suppress the transcription of *eef1a1l2* and speculated that Nanog safeguards oocyte maturation by suppressing *eef1a1l2* during oocyte development in zebrafish. However, zebrafish Nanog is well-known for acting as a transcription activator in zygotic genome activation (ZGA) and shaping the embryo during gastrulation. For instance, together with Pou5f3 and SoxB1, Nanog initiates the zygotic gene activation (ZGA), of which miR-430 is directly activated by Nanog and responsible for clearance of maternal mRNA during MZT (Lee et al., 2013). Further studies proved Nanog binds to the high nucleosome affinity regions (HNARs) center and opens chromatin with Pou5f3 and Sox19b synergistically, primes genes for activity during ZGA in zebrafish (Veil et al., 2019; Palfy et al., 2020). Nanog directly activates *mxtx2* and regulates the extraembryonic tissue and and embryonic architecture formation (Xu et al., 2012; Gagnon et al., 2018; Veil et al., 2018). Quantitative imaging also reveals Nanog cooperates with Pou5f3 to promote ventral fate (Perez-Camps et al., 2016). These studies illustrate that Nanog switches from transcriptional repressor to transcriptional activator during oocyte-to-zygote transition. As a homeodomain protein, Nanog binds with target genes at homeobox domain, while which repressive partner interacts with Nanog to exhibit gene silence function in oocyte still needs to be further investigated.

## Methods and Materials

### Zebrafish maintenance

All the zebrafish used in this study were maintained and raised as previously described (Westerfield, 1995) at the China Zebrafish Resource Center of the National Aquatic Biological Resource Center (CZRC-NABRC, Wuhan, China, http://zfish.cn). The Wildtype (WT) embryos were collected by natural spawning from AB strain. Oocyte developmental stages were classified according to previous studies (Selman et al., 1993; Lubzens et al., 2010). Developmental stages of mutant embryos were indirectly determined by observation of WT embryos born at the same time and incubated under identical conditions. The experiments involving zebrafish were performed under the approval of the Institutional Animal Care and Use Committee of the Institute of Hydrobiology, Chinese Academy of Sciences under protocol number IHB2014-006.

### Double mutant generation of *nanog* and *eef1a1l2*

M*nanog* was generated by crossing *nanog*^-/-^ female with WT male as previously described (He et al., 2015; He et al., 2020). *eef1a1l2* mutant was generated in *nanog* mutant background using CRISPR/Cas9. The gRNA target and PAM sequence (underlined) of *eef1a1l2* is 5’-GGCCACCTCATTTACAGTG<u>TGG</u>-3’, pT3TS-zCas9 was used for Cas9 mRNA transcription; capped Cas9 mRNA was generated using T3 mMessage Machine kit (AM1344, Ambion). gRNA was generated using in vitro transcription by T7 RNA polymerase (Promega, USA). Cas9 mRNA and gRNA were co-injected into embryos crossed by *nanog*^+/-^ female and *nanog*^-/-^ male at one-cell stage. M*nanog*/M*eef1a1l2* was obtained by crossing *nanog*^-/-^,*eef1a1l2*^-/-^ female with WT male. The primers used for mutant screening are listed in S2 Table.

### Morphological analysis of ovary and oocyte

After MS-222 anesthesia, we dissected the intact gonadal tissues from WT, *nanog*^-/-^, and *nanog*^-/-^,*eef1a1l2*^-/-^ adult zebrafish (4-months post fertilization) and calculated the gonadosomatic index (GSI; gonad weight/body weight X 100%). For embryos, chorion elevation distance and oocyte diameter were measured at 15 minute-post-fertilization (mpf) using ImageJ. Oocyte diameter was the longest distance in the vertical direction of the animal-vegetal axis. Chorion diameter was the longest length of chorion when it was fully inflated.

### Oocyte isolation, in vitro culture and germinal vesicle breakdown (GVBD) assay

Ovaries were dissected from adult females and transferred into oocyte sorting medium (OSM), made from 90% Leibovitz’s L-15 medium (Gibco, USA) and 10% Fetal Bovine Serum (BI, Israel) with 100μg/ml Penicillin-Streptomycin (Gibco, USA). Oocytes were manually separated and divided into five groups based on their size and vitellogenic state: primary growth stage (stage I), previtellogenic stage (stage II), vitellogenic stage (stage III), full-growth stage (stage IV) and the oocyte after GVBD in vitro was defined as stage V (mature oocyte). Ovulated maturation oocyte was defined as egg. Different stages of oocytes were gently separated using two tweezers in a dish covered with 1% agarose.

Dissociated stage IV oocytes were transferred into oocyte culture medium (OCM) by gentle pipetting. OCM was made from 90% Leibovitz’s L-15 medium and 10% Fetal Bovine Serum with 1 μg/mL 17α-20β-dihydroxy-4pregnen-3-one (DHP, Cayman, USA). Sorted oocytes were cultured at 28°C for 2h according to previous study (Nair et al., 2013). The GVBD rates were determined in a unified standard by ImageJ. The concentrations of different inhibitors added to OCM were *eif4a* inhibitor (25ng/μL, SantaCruz, USA), ISRIB (5μM, Selleck, Shanghai), GSK2606414 (50nM, Selleck, Shanghai).

### Cortical Granules (CGs) staining

Ovulated eggs at 10 minute-post-activation (mpa) in water were collected and fixed with 4% paraformaldehyde (PFA) overnight for further steps. Cortical granules were visualized by staining embryos with 50 μg/ml FITC-conjugated *Maclura pomifera* agglutinin (Vector Laboratories, FL-1341) as previously described (Mei et al., 2009).

### SDS-PAGE and Coomassie staining

SDS-PAGE and Coomassie staining were performed following the established protocol (Schagger, 2006). To obtain yolk protein, 10 oocytes were cleaved in 500 μL TNE buffer, made from 10 mM Tris-HCl (pH=7.4), 150 mM NaCl, 5 mM EDTA and 1% Triton X-100. 10 μL lysate was loaded for SDS-PAGE and Coomassie staining as previously described (Sun et al., 2018). Intensity measurement was done using ImageJ.

### Western blot analysis

GFP protein was purchased form DIA-AN Biotechnology and 20 pg of GFP protein per embryo was co-injected at one-cell stage. Injected embryos or dissected ovaries were homogenized using RIPA (P0013B, Beyotime). Western blot was carried out as previously described (Ye et al., 2019). Primary antibodies and dilutions for western blot were GAPDH (2058, DIA-AN, 1:3,000), mCherry (BE2026, Easybio, 1:2,000), GFP (2057, DIA-AN, 1:2,000), Hspa5 (11587-1-AP, Proteintech, 1:2,000), Ddit3 (AC532, Beyotime, 1:2,000), S6 (2217S, CST, 1:1,000), pS6 (2215S, CST, 1:1,000).

### RNA-seq and analysis

Total RNA of ovulated eggs of WT and *nanog* homozygous were extracted using TRIzol Reagent (Invitrogen) and mRNA was enriched using oligo-dT magnetic beads. First-strand cDNAs (from purified mRNA) was synthesized using random hexamers. The PCR-amplified cDNA was purified using AMPure XP beads, then 1□μL cDNA was validated using an Agilent 2100 Bioanalyzer. Sequencing libraries were generated using the Illumina TruSeq RNA sample preparation kit v2 according to the manufacturer’s recommendations. Clustered library preparations were sequenced on an Illumina HiSeq^TM^ 2000 machine and 100 bp single-end reads were generated. Clean reads, with low quality reads removed from the raw data, were mapped to the zebrafish GRCz10 reference genome using TopHat2 (Kim et al., 2013). HTSeq v0.6.1 was used to count the read numbers mapped to each gene. Then, the RPKM of each gene was calculated to determine gene expression levels (Trapnell et al., 2010). Differential expression analysis was performed using the DESeq (Anders and Huber, 2010). Genes with an adjusted P-value□<□0.05 as calculated by DESeq were considered differentially expressed. The original RNA-seq data has been deposited to the BioProject with accession number PRJNA633216.

### Proteomics

Ovulated eggs of WT and *nanog* homozygous were pooled and homogenized for quantitative proteomic analysis. The iTRAQ analysis was performed as previously described (Miao et al., 2017). The UniPort proteome sequence for Danio rerio were used for the database searching.

### Chromosome-immunoprecipitation PCR (ChIP-PCR)

Chromatin immunoprecipitation (ChIP) assays were performed with a ChIP assay kit (Cell Signaling Technology) as described (Wei et al., 2014). Briefly, 2 ovaries of *Tg*(CMV:*nanog-myc*) at 6mpf were dissected and lysed for ChIP assay. Immunoprecipitation was carried out using anti-Myc antibody (CST). The immunoprecipitation of *eef1a1l2* genomic in immunoprecipitated chromatin was detected by PCR. primers specific for the *eef1a1l2* promoter region were used and the sequences were listed in S2 Table. The exon of ribosomal protein *rpl5b* was served as a negative control, and the primers were 5′-GGGGATGAGTTCAATGTGGAG-3′ (forward) and 5′-CGAACACCTTATTGCCAGTAG-3′ (reverse) as described (Belting et al., 2011).

### Transmission electron microscopy (TEM) analysis

Isolated early-stage oocytes from different genotypic ovaries were collected into a test tube and fixed with 100 μL 2.5% glutaraldehyde at 4°C overnight. Samples preparation for transmission electron microscopy (TEM) was according to the previous described (Zhang et al., 2021) and observed under TEM (Hitachi, HT7700).

### In situ hybridization

PCR-amplified sequences of genes of interest were used as templates for the synthesis of an antisense RNA probe, labelled with digoxigenin-linked nucleotides. Whole-mount *In situ* hybridization (WISH) on embryos were performed as described previously (Thisse and Thisse, 2008). For *in situ* hybridization on frozen section, adult ovaries were stripped and embedded in Optimal Cutting Temperature compound (O.C.T. SAKURA) and sectioned at 10 μm. The procedures of hybridization followed a previous study (Zhang et al., 2020).

### Immunofluorescence

For whole-mount immunofluorescence, embryos were collected and fixed in 4% PFA for overnight at 4°C. For immunofluorescence on cryosections, sections were prepared as in situ hybridization and fixed in 4% PFA for 20 min at room temperature. The procedure of immunostaining on embryos and slides are referred to previous studies, separately (He et al., 2020; Zhang et al., 2021). Primary antibodies and dilutions were Cleaved-Caspase3 (#9661, CST, 1:1,000), Myc (#2276, CST, 1:1,000).

### Reverse-transcription quantitative PCR (RT-qPCR)

Total RNA was extracted from indicated samples using TRIzol method. The RNA was reverse-transcribed with PrimeScript™ RT reagent Kit (Thermo) and relative abundance of target mRNAs was examined with gene-specific primers. *gapdh* was used as a normalization control. Sequences of PCR primers are listed in S2 Table. RT-qPCR was performed using the SYBRGreen Super mix from BioRad (USA) on a BioRad CFX96.

### Stem-loop RT-PCR of miR-430a

Stem-loop RT-PCR was performed to quantify the expression of miR-430a as previously described (Chen et al., 2005). Total RNAs were reversely transcribed using the miR-430a specific primers and U6 was used as internal control. The PCR primers of miR-430a and U6 used in this study referred to previous study (He et al., 2020).

### Terminal deoxynucleotidyl transferase dUTP nick end labeling (TUNEL) assay

Ovaries were dissected from 4 months post fertilization WT and *nanog*^-/-^ adult fish and cryosections were prepared as in situ hybridization. The samples were sectioned at 10 μm thickness. The TUNEL cell death assay was performed using the In Situ Cell Death Detection Kit (Roche) according to the manufacturer’s instructions. Images were obtained using a laser scanning confocal microscope (Leica SP8).

### Statistical analysis

GraphPad Prism 8.3.0 software was used for statistical analyses and statistical graphs. Significance of differences between means was analyzed using Student’s t test. Sample sizes were indicated in the figure legends. Data were shown as mean ± SD. *P* value below 0.05 marked as *, *P* value below 0.01 marked as **, and *P* value below 0.001 marked as ***, ns means no significant difference.

## Acknowledgements

We thank other members of the Sun laboratory for discussions, and Linglu Li from the China Zebrafish Resource Center for fish raising assistance.

## Data availability

RNA-seq data in this study have been deposited in SRA under the accession number PRJNA633216. Raw data of the iTRAQ proteome analyses are available in iProX under the accession number IPX0004874000.

